# Numerical validation of 3D-VAR data assimilation for estimating network dynamics in multivariate EEGs

**DOI:** 10.1101/2025.05.22.655248

**Authors:** Hiroshi Yokoyama, Keiichi Kitajo

## Abstract

Data assimilation (DA) is widely used as a data-driven system identification method, enhancing our understanding of the generative mechanisms in the observational data based on the identified model and its parameters. Due to this advantage, DA has gradually begun to be applied in neuroscience studies. Recently, we introduced a DA-based method for reconstructing the internal state of the brain using the neural mass model and human scalp electroencephalography (EEG). This method allows us to estimate the balance of synaptic interactions between excitatory and inhibitory neurons (E/I balance) in the brain only from observed EEGs. Although we confirmed the neurophysiological validity of the proposed EEG-DA method in our recent works, this method cannot be applied to multivariate EEGs to parallelly infer the E/I balance changes and underlying network dynamics. In this study, to address this issue, we proposed an extended version of our DA method specified to multivariate observed EEGs. This method enables us to estimate sensor-level functional network and E/I balance changes of each EEG sensor in parallel. The method was validated by showing that it could parallelly estimate sensor-level network structures and E/I balance changes from synthetic multivariate EEGs. The results of this study indicate that it has the potential to quantify how the E/I balance changes functionally affect the brain network dynamics.

## 1 Introduction

Data assimilation (DA) is widely used as a data-driven system identification method in practical applications such as weather forecasting [21, 23]. Recently, it has begun to be applied in various research fields, including neuroscience [3,14,20,25,28]. The advantages of applying DA for neuroscience studies include not only enabling accurate time-series forecasting in neual data, but also enhancing our understanding of the generative mechanisms in the observed neural data based on the identified systems and their parameters.

For this reason, we recently developed the DA-based method to estimate the latent state of the brain from human scalp electroencephalographic signals (EEGs) [31,32]. Our focus of this method is on estimating the balance of synaptic interactions between excitatory and inhibitory neural ensembles (E/I balance) only from EEGs. In this method, we use a noise-adaptive Ensemble Kalman Filter (EnKF) [7, 26, 29, 31, 32] to directly fit the observed EEG data to the neural mass (NM) model known as a computational model of EEGs. Applying our proposed EEG-DA method, we succeeded in reconstructing the time-varying changes in the parameters of the NM model related to the dynamics of the E/I balance at the sub-second scale with non-invasive measurements (i.e., EEGs). Since the disruption of E/I balance in the brain can lead to various neuropsychiatric disorders [2, 8, 9, 13, 16, 18, 19, 22, 30], our EEG-DA method could enhance our understanding of the functional roles of these changes. Furthermore, the neurophysiological validity of E/I balance estimation in our method was also confirmed in our recent works [31, 32].

However, our EEG-DA method can be applied to only a single time series of EEGs. Therefore, this method cannot rigorously estimate E/I balance changes underlying whole brain network dynamics. To address this limitation, we extend our previous EEG-DA method to enable the simultaneous estimation of E/I balance and functional brain network. In our new EEG-DA methods, the sensor-level activities in the multivariate observed EEGs are directly assimilated into the network neural mass (NNM) models with three-dimensional variational (3D-VAR) DA algorithms [1, 17, 20, 21, 27]. Moreover, to consider the observation noise effect for system identifications, we applied a combined method of 3D-VAR with data-driven observation error covariance tuning [4, 6], enabling noise-adaptive state estimation in the NNM model.

In this study, to confirm the numerical validity of our proposed method, we applied the method to synthetic multivariate EEGs generated by the NMM model with known E/I parameters and network structures. By doing so, we confirmed that the estimated model parameters and network structures were consistent with the true ones.

The organization of this manuscript is as follows. We first introduce our proposed method in Section 2. We provide the simulation procedures and their results for numerical validations of our EEG-DA method in the later part of Section 2 and Section 3. Conclusions are given in Section 4.

## 2 Materials and Methods

We first explain the basics of NNM model in Section 2.1. The mathematical details for two key methods in our implemented DA algorithm: (1) 3D-VAR [17, 27] and (2) error covariance tuning [4, 6] are provided in Sections 2.2 and 2.3. In Section 2.4, we describe the procedures of numerical simulation to confirm the validity of our implemented EEG-DA method.

### 2.1 Descriptions of network neural mass model

In this section, we will provide the description of NNM model that is applied to the physical model of multivariate EEGs in our proposed DA method.

As mentioned in Introduction section, we recently proposed the EEG-DA method by applying the NM model to the single time-series of EEG [11, 31, 32]. In the current study, to extend our EEG-DA proposed in recent works [31, 32] for multivariate EEG data, we applied the network of NM models (i.e., NNM model) [10, 12] as the state model of EEG observations. We assumed that the network dynamics of EEGs formulated as the interactions between pyramidal cells, excitatory, and inhibitory interneurons in the NNM model [10, 12]. In our implemented method, the dynamics for each sensor of EEGs correspond to each functional node of the NNM model, and sensor-level interactions of EEGs are considered network interactions in NNM models. The NNM models for each functional node are formulated as the following differential equations:

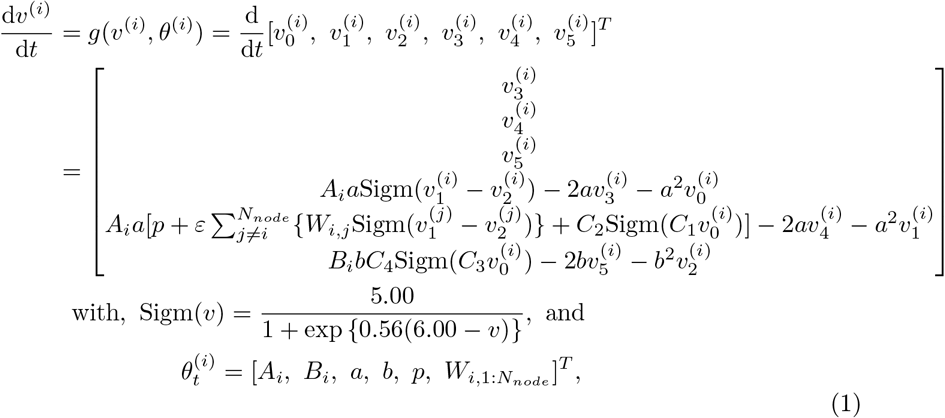

where variables 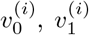, and 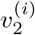 indicate the mass of postsynaptic potential (PSP) in pyramidal, excitatory, and inhibitory neural populations. The subscripts *i* of the variable *v* corresponds to the index of node number and time stamp, respectively. *N*_*node*_ stands for the maximum number of the node. The simulated EEGs in the *i*-th node of the NNM model are defined as 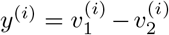. The parameters, *A*_*i*_, *B*_*i*_, *a, b*, and *p*, are the target-to-state estimation in our proposed EEG-DA method. *A*_*i*_ and *B*_*i*_ indicate the excitatory and inhibitory synaptic gains, which control the amplitude of the excitatory PSP (EPSP) and inhibitory PSP (IPSP) in the *i*-th node. *W*_*i,j*_ stands for the network coupling strength for each pair of nodes in the NNM model. Other parameters *a, b*, and *p* indicate the inverse of the time constant for E/I neurons and the background noise input of the model. In our proposed EEG-DA method, while parameters *A*_*i*_, *B*_*i*_, *W*_*i,j*_ are estimated for each node or pair of nodes, other parameters *a, b* and *p* are estimated as the common parameters in each network node. The fixed constants *C*_*n*_ (where, *n* = 1, …, 4) account for the number of synapses. *C*_*n*_ was fixed as *C*_1_ = 135, *C*_2_ = 108, *C*_3_ and *C*_4_ = 33.75, based on previous studies [5, 11]. *ε* indicates the global coupling strength that is set as *ε* = 1.0 in this study based on the ref [12].

To apply the NNM model to our EEG-DA method, the Eq. (1) were transformed to the state-space form as follows:

state model;

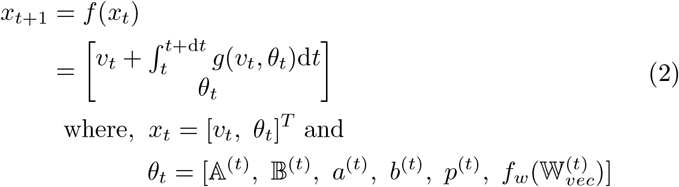

observation model;

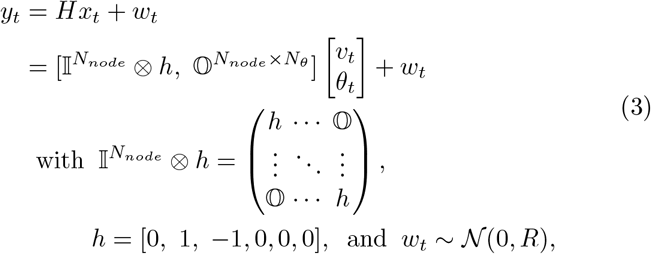

where, 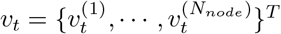 and *θ*_*t*_ indicate the state variables and parameters of the NNM model, respectively. Note that 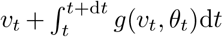 indicating numerical integral of the NNM model was implemented with the fourth-order Runge-Kutta method with time step d*t* (sampling interval). 𝕎 _*vec*_ = vec(*W*_*\{i*=*j}*_) indicates the vectorized variables containing the off-diagonal elements of network coupling strength 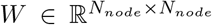. 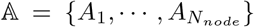 and 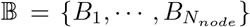 indicate the vector of excitatory and inhibitory gain parameters in each node, respectively. Other variables, *a, b*, and *p*, are set as common parameters for all nodes in the NNM model. In the state model for the model parameter *θ*_*t*_, to set the boundary constraint for network coupling strength with 0 ≤ *W* ≤ 1, the state model of this parameter is defined as 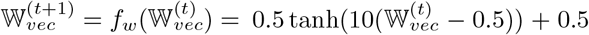. Other parameters defined as constants such as 𝔸 ^(*t*+1)^ = 𝔸 ^(*t*)^. By applying the above state-space form of the NNM model to the 3D-VAR DA method, the state and parameters in this model are estimated in a data-driven manner.

### 2.2 Model identification with noise adaptive 3D-VAR

In this section, we introduce the mathematical details of our implemented noise adaptive 3D-VAR algorithm and procedures for applying this algorithm to the EEG-DA. The optimal state *x* in the state-space form of the NNM model can be solved to maximize the following posterior likelihood.

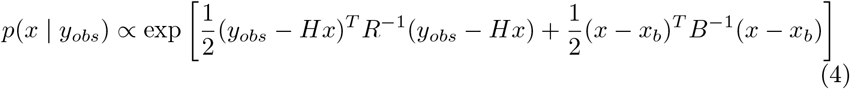

From the viewpoint of the Bayesian theorem, maximizing the above likelihood can be considered as minimizing the following cost function 𝒥 (*x*) [17, 27].

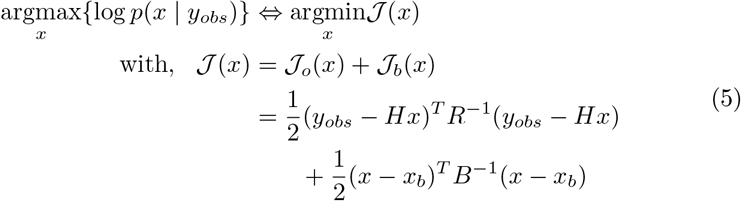

In this case, the above problem could be analytically solved as below [17, 27],

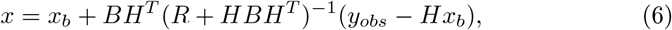

where *B* and *R* indicate background error (state error) and observation error covariance matrices, respectively. *y*_*obs*_ and *x*_*b*_ are observation and background state variables, respectively. The background state *x*_*b*_ refers to the preliminary or prior estimated value of the state *x*. When applying the 3D-VAR algorithm to estimate the state *x*, both error covariances *B* and *R* should be well-specified as known parameters; however, both *B* and *R* are typically unknown. Therefore, in an ideal manner, both *B* and *R* should be estimated in a data-driven manner.

To estimate *B* and *R* optimally, we now assume that these error covariance matrices can be iteratively deduced by applying the tuning parameters *s*_*b*_ and *s*_*R*_ as below [4, 6].

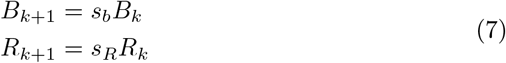

Note that *B*_*k*_ and *R*_*k*_ indicate the estimated covariances in *k*-th iteration. For estimating *B* and *R* iteratively with the Eq. (7), the tuning parameters *s*_*b*_ and *s*_*R*_ should be determined; however, these parameters are unknown. In this case, assuming that the optimal set of error covariance (*B, R*) are determined under the minimization of cost function *𝒥* (*δx*), the following equalities would be satisfied as demonstrated by previous works [4, 6].

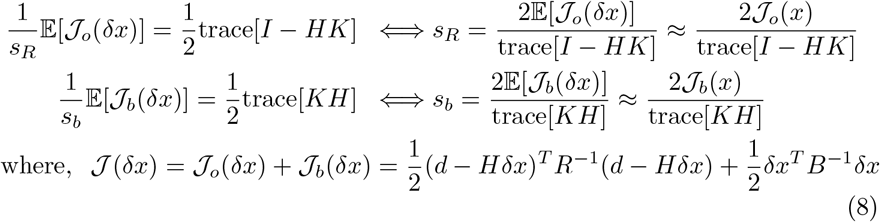

Note that *δx* = *x* − *x*_*b*_ and *d* = *y*_*obs*_ − *Hx*_*b*_ indicate the background and observation error. Based on the above equations, the analytical solution in the parameters *s*_*b*_ and *s*_*R*_ would be given, using the estimated state *x* in Eq. (6). Therefore, applying the resulting values of *s*_*b*_ and *s*_*R*_ in Eq.(8), both error covariance *B* and *R* can be determined by using Eq.(7) in a data-driven manner.

### 2.3 Implemented algorithms

In the prior sections, we explained the mathematics regarding two key methods for our implemented EEG-DA algorithms: 3D-VAR [17, 27] and error covariance tuning [4, 6]. In the following section, we will briefly provide the implemented algorithm and procedures for data-driven identification of the NNM model with observed EEGs. The outline of our implementations is shown in Fig. 1.

**Fig. 1.**
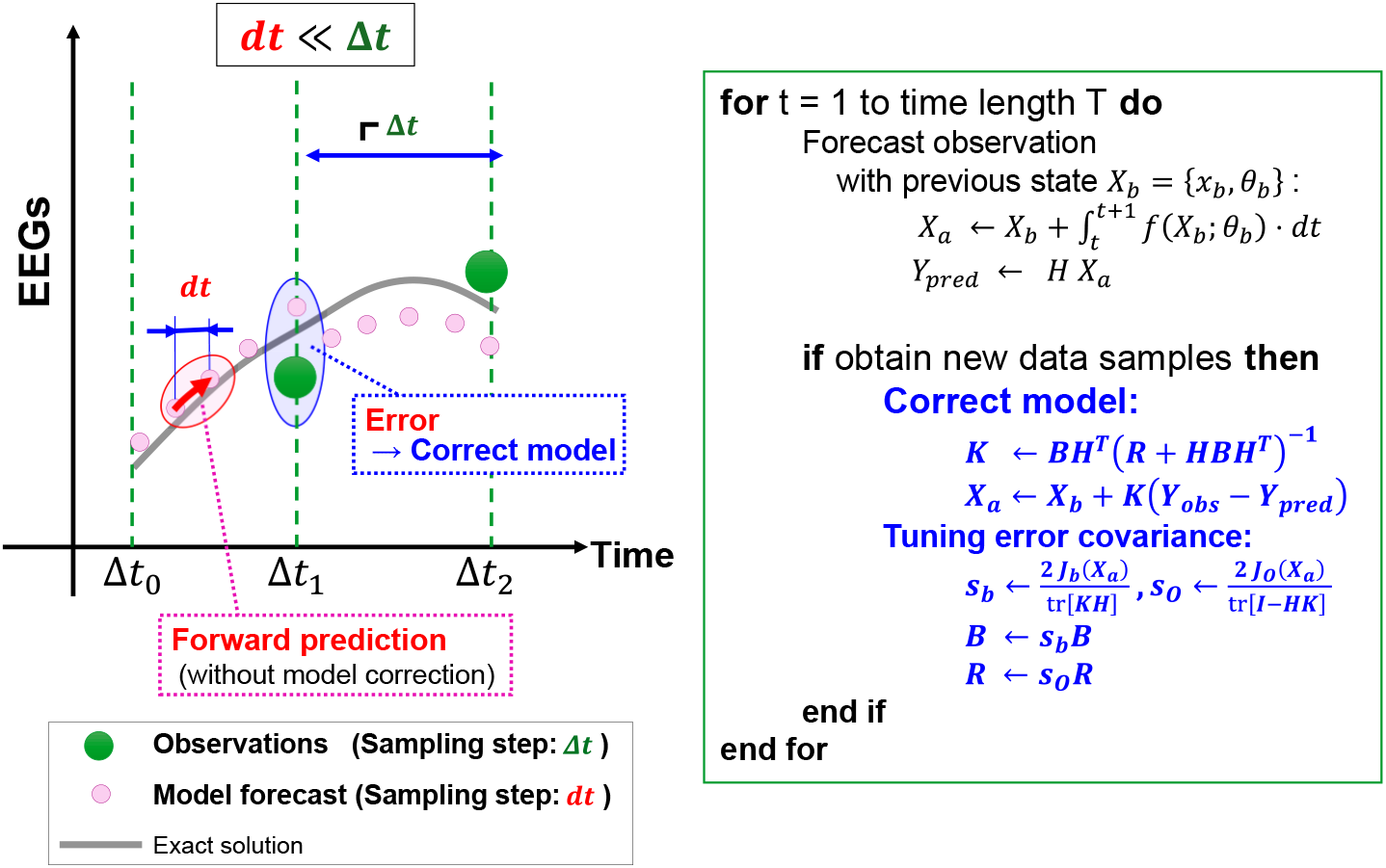
Schematic illustration and pseudo-code of model identification procedures in our EEG-DA algorithm.

As illustrated in Fig. 1, our EEG-DA consists of two steps: (1) forward prediction and (2) model correction. In the forward prediction step, the time-series prediction of multivariate EEGs is obtained from the time integration of the NNM model with a time step d*t* based on the fourth order Runge-Kutta method as well as the state model in the Eq.(2). For this prediction, the model state *v*_*t*_ and parameters *θ*_*t*_ in the NNM model (see Eqs. (1 and 2)) are set as an initially given or priorly estimated values. If we can obtain the EEG observations, the model correction step is applied after calculating the forward predictions of EEGs. Through this procedure, the model state and parameters in the NNM model are corrected to minimize the EEG prediction error based on the noise adaptive 3D-VAR mthod with Eqs. (6–8), described in Section 2.2. Moreover, in our implementations, we assumed that the EEG observations would be sparsely sampled relative to the time step of the model prediction [1]. Therefore, the time-step of observation sampling is set as *Δt* which is a larger size of time-steps relative to model prediction d*t* (i.e., *dt* ≪ *Δt*). For example, if we set the time-step of model prediction (pink-dot in Fig. 1) and observations (green-dot in Fig. 1) as *dt* = 0.0005 (i.e., sampling frequency: 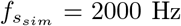) and *Δt* = 0.0025 (i.e.,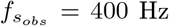) respectively, the observed data is obtained at every 5th time step in the model predictions [1]. These steps for our implemented EEG-DA are applied to the whole time intervals of the target observational data. By doing so, the model state and parameters in the NNM model are optimally estimated. The algorithms in these procedures are based on the combined algorithm with the 3D-VAR and observation error covariance tuning scheme. The pseudo-code of the above procedures are also shown in Fig.1 with schematic illustration.

### 2.4 Numerical simulation: twin experiment

#### Outline and main aim of the simulation

To confirm the validity of our proposed method, we conducted the numerical simulation based on the twin experiment framework. In this simulation, we applied our EEG-DA method to synthetic EEG data generated by the five-node NNM model with known parameter settings and network structures. By applying our EEG-DA method to such generated synthetic EEGs, we tested whether our method could correctly reconstruct the network structures and the parameters associated with the dynamics of E/I balance in the NNM models from synthetic data. The simulation was executed 50 times with varying initial conditions, while consistently keeping the parameters and network structures. This approach allowed us to evaluate the robustness and accuracy of the EEG-DA method by averaging the performance over these 50 estimations.

Moreover, in these numerical simulations, we also focused on the following two issues: (i) the effect of the sample length of the observations and (ii) the impact of sampling intervals for the observations. For the first issue, we aimed to reveal the effect of the total sample number for the system identifications in our 3D-VAR-based EEG-DA. To do so, the generated time-series of synthetic EEGs were divided into two time periods: training period *T*_*train*_ and test period *T*_*test*_, before applying these data to our EEG-DA method. The time length of *T*_*train*_ period was selected from *T*_*train*_ = 20, 40, and 60 (seconds). The synthetic EEGs in *T*_*train*_ were applied to our EEG-DA method for model identifications (i.e., parameter estimation). After identifying the model in *T*_*train*_ period, the prediction accuracy of EEG observations was evaluated in the test period *T*_*test*_ by using the identified model in *T*_*train*_. The time length of *T*_*test*_ was fixed as *T*_*test*_ = 20 (seconds). For the second issue, we also focused on the sensitivity of prediction accuracy relative to the intervals *Δt* for the observation sampling in the numerical simulation. As shown in Fig. 1, we assumed that observed EEGs were sparsely sampled with *Δt* when applying them as the observed data in our 3D-VAR-based EEG-DA scheme. The sampling interval *Δt* of observed EEG is selected to satisfy the relationship *Δt* ≪ d*t*, where d*t* indicates the interval of the model prediction in the 3D-VAR scheme (Fig.1). In the whole simulations, time intervals d*t* are set as *dt* = 0.0005 (i.e., the sampling frequency is fixed as 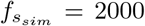). Meanwhile, *Δt* (i.e., the interval of the observation sampling) is selected from *Δt* = 0.001, 0.01, and 0.2 (i.e., sampling frequency: 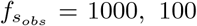, and 5 Hz) to consider the effect of the differences between *Δt* and d*t*. Thus, in this evaluation, we considered the sensitivity of model prediction accuracy under the condition for the ratio of sampling frequency between the model prediction and observation sampling with 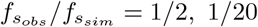, and 1*/*4000.

We note that the synthetic EEG was generated with the same time-step interval *dt* = 0.005 (i.e.,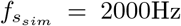) of model prediction in our EEG-DA scheme, because of the accuracy evaluation for the EEG predictions in *T*_*test*_ period. Therefore, while the synthetic EEGs were generated under the same time steps d*t* in the model predictions on our EEG-DA scheme, these synthetic EEGs were sparsely sampled in the *T*_*train*_ period with time-step *Δt* when applying these EEGs as the observations for our implemented 3D-VAR algorithm.

#### Generation procedures of synthetic EEGs

As mentioned above, the synthetic EEGs were generated by the five-node NNM model using fourth-order Runge-Kutta time integrals with time-step *dt* = 0.005 time-steps (i.e., sampling frequency 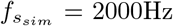). The parameter setting of synthetic EEGs is set as below. The true parameter of network structures 𝕎 is selected so that network structures follow as in Fig.2B (see “true” network diagram in this figure). The weight of all edges is set as *W*_*i,j*_ = 0.9 if a connection exists from *j*-th to *i*-th node. Other parameters for each network node *i* were set as standard parameters to produce alpha-like oscillations (E/I synaptic gain parameters: *A*_*i*_ = 3.25 and *B*_*i*_ = 22.0, E/I time decay parameters: *a*_*i*_ = 100 and *b*_*i*_ = 50, external input *p*_*i*_ ∼ *𝒩* (220, 20)), based on the studies [5, 11]. Furthermore, the E/I synaptic gains for each node *i, A*_*i*_ and *B*_*i*_, are added perturbations *n*_*i*_ that followed *n*_*i*_ ∼ *𝒩* (0, 0.01). Based on these settings, the synthetic data were generated 50 times with different initial values of model state *v*_0_ in the NNM models for each simulation condition. After generating the synthetic EEGs based on the above parameter settings, to prepare the noise-distorted observations, the Gaussian noise satisfying the noise scale with 10 dB of signal-to-noise ratio was added to all generated synthetic EEGs. After treating all the above procedures, the synthetic EEGs were applied to the twin experiment test as described in the prior section.

**Fig. 2.**
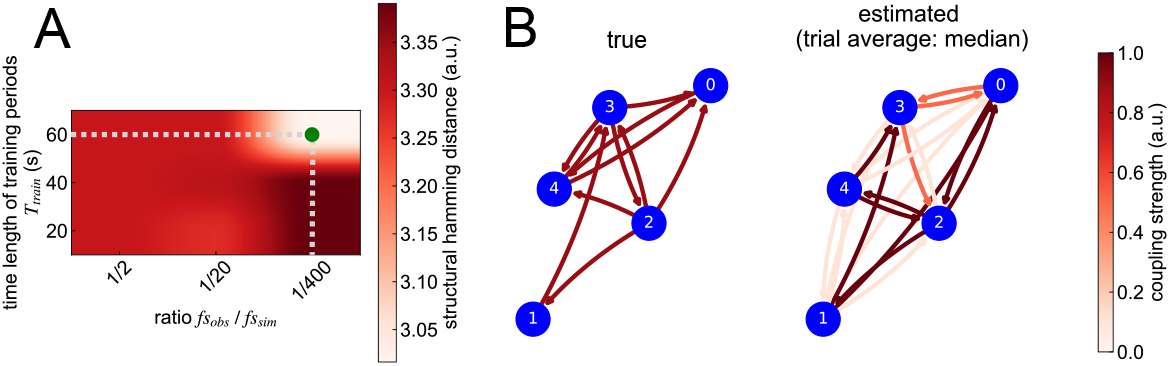
Sensitivity of network parameter reconstructions in our 3D-VAR based EEG-DA. (A) Sensitivity in the estimation of the network parameter *W* relative to time-length *T*_*train*_ and ratio 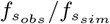. The green dot indicates the most accurate condition. (B) Comparisons of estimated network structures with true network structures. The estimated network structure was obtained from a median of estimated *W* over 50 repetitions under the most accurate condition in (A).

#### Evaluation index of estimation

To quantify the prediction performance in our EEG-DA method, we applied three metrics: structural Hamming distance (SHD), mean absolute error (MAE) [31], and phase coherence (PC) [15]. SHD [24] was applied to calculate the dissimilarity between true and estimated network coupling strength. For the evaluation of time-series prediction accuracy, we applied two indexes: MAE to assess the prediction error of EEG amplitude, and PC to evaluate the prediction accuracy of EEG phase.

## 3 Results

In this study, we proposed a 3D-VAR-based EEG-DA scheme for estimating E/I balance changes and network structures simultaneously. To confirm the numerical validity of this method, we conduct a twin experiment-based simulation. As a result, we successfully obtained the preliminary findings suggesting that our method can be applied to practical EEG data to interpret the functional role in E/I balance changes underlying whole-brain network dynamics. In this section, we will explain these results and discuss them.

### 3.1 Network reconstruction

We first demonstrate the parameter reconstruction accuracy obtained from the results through 50 repetitive simulations (Fig.2 and Table 1). Fig.2 showed the network estimation results. As shown in Fig.2A, even though we did not find the significant differences in the SHD score depending on both conditions: time length *T*_*train*_ and ratio of sampling frequency 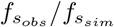, the smallest SHD score was found under the conditions: *T*_*train*_ = 60(sec) and the ratio: 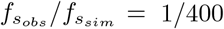 (i.e., the observed EEGs are sampled at every 400th time-step in the model prediction). Since the SHD score indicates the dissimilarity of the estimated network structure with the true one, the condition with the smallest SHD means that the most accurate estimation is obtained in this condition. As can be seen in Fig.2B, the averaged estimation of network structure in this condition was quite similar to the true network. Furthermore, it was observed that the E/I synaptic gain parameters (*A*_*i*_, *B*_*i*_) were precisely determined under similar conditions, which resulted in the most precise network estimations (Table 1).

**Table 1.** Estimation of E/I synaptic gain parameters. *A*_*i*_ and *B*_*i*_ indicated the E/I synaptic gain parameters for each node *i* under the conditions *T*_*train*_ = 60 (sec) and 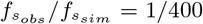. Each estimated values were listed with mean *±* 3 *×* standard deviation obtained from 50 times estimation results.

**Excitatory gain parameters**
| | $A_0$ | $A_1$ | $A_2$ | $A_3$ | $A_4$ |
| --- | --- | --- | --- | --- | --- |
| true | 3.170 | 3.282 | 3.247 | 3.314 | 3.220 |
| prediction | $3.248 \pm 0.037$ | $3.253 \pm 0.037$ | $3.250 \pm 0.024$ | $3.251 \pm 0.036$ | $3.249 \pm 0.027$ |

**Inhibitory gain parameters**
| | $B_0$ | $B_1$ | $B_2$ | $B_3$ | $B_4$ |
| --- | --- | --- | --- | --- | --- |
| true | 22.04 | 21.99 | 21.95 | 22.06 | 21.95 |
| prediction | $22.00 \pm 0.024$ | $22.00 \pm 0.031$ | $22.00 \pm 0.035$ | $22.00 \pm 0.031$ | $22.00 \pm 0.027$ |

### 3.2 Prediction accuracy in EEG observations

Finally, we showed the prediction accuracy of observed EEGs (Fig.3). While Fig. 3A demonstrated the sensitivity of the EEG amplitude estimation relative to the changes in *T*_*train*_ and ratio 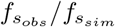, Fig.3B is those of the EEG phase estimation. Moreover, these sensitivity analyses were compared with the different 2-second intervals in *T*_*test*_ period: early (0.0 − 2.0 sec), middle (8.0 − 10.0 sec), and later (18.0 − 20.0 sec).

**Fig. 3.**
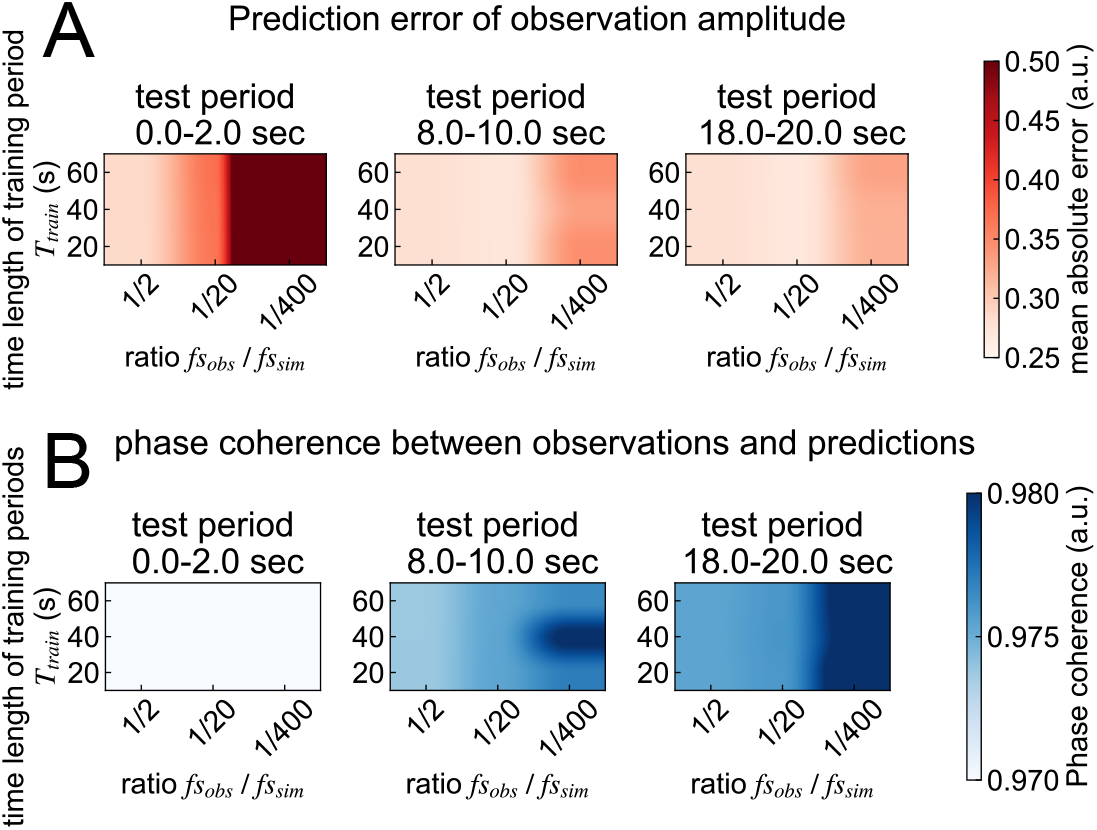
Sensitivity of EEG time-series predictions in our 3D-VAR based EEG-DA. (A) Results of sensitivity in prediction error of EEG amplitudes, relative to the changes in time-length *T*_*train*_ and ratio of the sampling frequency between observations and model predictions 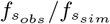. The color bar displays the MAE, indicating the higher values are higher errors. (B) Results of sensitivity in prediction error of EEG phase, relative to the changes in *T*_*train*_ and 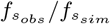. The color bar displays the phase coherence, indicating the higher values are smaller phase estimation error.

As shown in Fig.3, the contrastingly opposite tendencies were obtained from sensitivity analysis between EEG amplitude and EEG phase estimation. The prediction error of EEG amplitude tends to be higher relative to the decrease in sampling frequency of the observations (Fig.3A). However, the prediction error of the EEG phase tends to be smaller relative to the reduction of the sampling frequency of the observations, because the phase coherence between observations and the prediction of EEGs would be higher relative to the reduction of the sampling frequency of the observations (Fig.3B).

Based on these tendencies, although sampling the observations at shorter intervals would lead to fine model identification achieving an accurate prediction of the amplitude in the immediate future state, the prediction ability for long-term periods could not be capable. In contrast, the sparse sampling for observations could not accurately estimate the amplitude; however, phase estimation would be accurate even in relatively long-term forecasts. The lower error of phase prediction indicates that the identified model in our EEG-DA algorithm could estimate the event timing of EEG dynamics with a tighter time-scale order. Since enabling the estimation of the timing-dependent dynamics in EEGs was more important to understand the neurophysiological mechanisms, the sparse sampling for observations in 3D-VAR would be preferred to establish the EEG-DA framework.

We showed the typical example of EEG time-series prediction obtained from one single trial under the condition: *T*_*train*_ = 60(sec) and 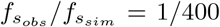(Fig.4). As shown in this figure, based on the identified model using our EEG-DA scheme during the *T*_*train*_ period, the prediction of the models can correctly track the trend of the next 10–15 seconds of the true EEG signal without model correction based on the observation.

**Fig. 4.**
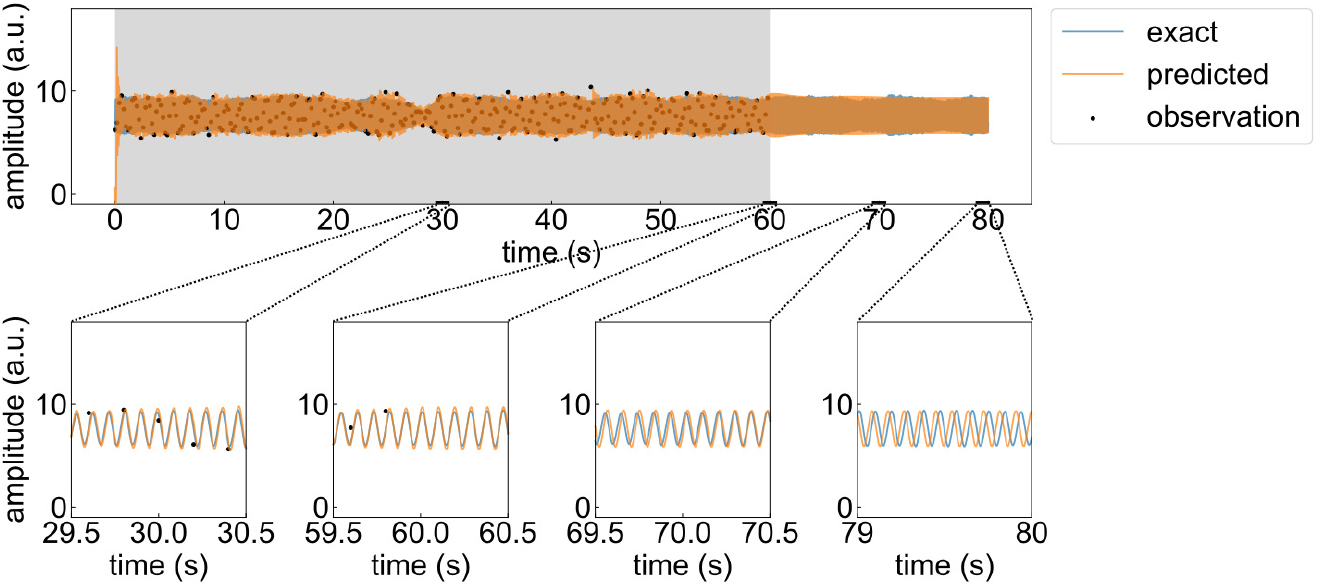
Example of EEG time-series prediction. Comparison between the observation and typical single trial prediction of EEG at the 4-th node under the condition: *T*_*train*_ = 60(sec) and 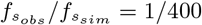. The upper panel shows the whole time series of EEGs. Gray shaded intervals indicates the training period *T*_*train*_ for our EEG-DA scheme. The interval between 60 and 80 s is test period *T*_*test*_. The lower four panels are enlarged views of the prediction results for each time period.

## 4 Conclusions

In this study, we proposed a noise adaptive 3D-VAR-based EEG-DA approach to simultaneously estimate the sensor-level network interaction and model parameters related to the E/I balance changes from observed multivariate EEGs. Through numerical simulations, our findings indicate the following two possibilities: (1) both network structures and E/I balance could be determined from EEGs using our implemented noise adaptive 3D-VAR scheme, (2) the identified model obtained by the noise adaptive 3D-VAR scheme could be applied to simulate the future state of the brain for the next few seconds. Indeed, the results of this study are just preliminary evidence. However, we think that the reliability of the estimation for E/I parameters and network structures was supported by the results obtained from 50 times repetitive estimations using the different sets of synthetic EEGs. Since the set of synthetic EEGs was generated by the NNM model with varying initial states while keeping the parameter settings, the simulation results in this study supported that our EEG-DA method can estimate the latent state of observational EEGs, independent of the initial state effects in dynamic systems. Nevertheless, the lack of neurophysiological validation for our EEG-DA method is an obvious limitation in this study. In our future work, we will focus on validating the neurophysiological aspects of our method by using the practically observed EEGs obtained from human patients.

## Acknowledgment

This research was partially funded by the JSPS KAKENHI grant (24K22305) from the Japan Society for the Promotion of Science, and the AMED Multidisciplinary Frontier Brain and Neuroscience Discoveries (24018835) from the Japan Agency for Medical Research and Development.

